# Mapping the risk of Rift Valley fever in Uganda using national seroprevalence data from cattle, sheep and goats

**DOI:** 10.1101/2022.05.12.491594

**Authors:** Dan Tumusiime, Emmanuel Isingoma, Optato B. Tashoroora, Deo B. Ndumu, Milton Bahati, Noelina Nantima, Denis Rwabiita Mugizi, Christine Jost, Bernard Bett

**Affiliations:** Ministry of Agriculture, Animal Industry and Fisheries, P.O Box 103, Entebbe, Uganda; International Livestock Research Institute, P. O. Box 30709-00100, Kampala, Uganda; United States Agency for International Development’s Bureau for Humanitarian Assistance 14 (USAID/BHA), Washington, DC; Global Health Support Initiative III, Social Solutions International, Washington DC; International Livestock Research Institute, P. O. Box 30709-00100, Nairobi, Kenya

**Keywords:** Rift Valley fever, seroprevalence, risk map, INLA, SPDE

## Abstract

Uganda has had repeated outbreaks of Rift Valley fever (RVF) since March 2016 when human and livestock cases were reported in Kabale after a long interval. The disease has a complex and poorly described transmission patterns which involves several mosquito vectors and mammalian hosts (including humans). We conducted a national serosurvey in livestock to determine RVF virus (RVFV) seroprevalence, risk factors, and to develop a risk map that could be used to guide risk-based surveillance and control measures. A total of 3,281 animals from 189 herds were sampled. Serum samples collected were screened at the National Animal Disease Diagnostics and Epidemiology Unit (NADDEC) using a competition multispecies anti-RVF IgG ELISA kit. Data obtained were analyzed using a Bayesian model that utilizes integrated nested Laplace approximation (INLA) and stochastic partial differential equation (SPDE) approaches to estimate posterior distributions of model parameters, and account for spatial autocorrelation that was detected in the data based on Moran’s *I* test. Variables considered included animal level factors (age, sex, species) and multiple environmental data including meteorological factors, soil types, and altitude. A risk map was produced by projecting fitted (mean) values, from a final model that had environmental factors onto a spatial grid that covered the entire domain. The overall RVFV seroprevalence was 11.39% (95% confidence interval: 10.35 – 12.51%). Higher RVFV seroprevalences were observed in older animals compared to the young, and cattle compared to sheep and goats. RVFV seroprevalence was also higher in areas that had (i) lower precipitation seasonality, (ii) haplic planosols, and (iii) lower cattle density. The risk map generated demonstrated that RVF virus was endemic in several regions including those that have not reported clinical outbreaks in the northeastern part of the country. This work has improved our understanding on spatial distribution of RVFV risk in the country as well as RVF burden in livestock.

**Author summary:** Outbreaks of Rift Valley fever occur periodically in Uganda in livestock and humans. An Initial outbreak was reported in Kabale in March 2016 after a long quiescent period. Factors that trigger these outbreaks have not been fully described, but above-normal precipitation and flooding are known to be important causes. This paper presents a study that was conducted in the country to analyse RVF virus seroprevalence and risk factors for exposure in livestock. The study also mapped spatial distribution of RVFV seroprevalence. Data used in the study were generated from a national cross-sectional serosurvey that involved cattle, sheep and goats. Results obtained showed that the national RVFV seroprevalence was 11.39%, with 95% confidence interval of 10.35 – 12.51%. Cattle had higher RVFV seroprevalence compared to sheep and goats. Environmental factors that were associated with increased seroprevalence were low precipitation seasonality, haplic planosols, and low cattle density. Prevalence maps generated showed that the southwestern, central and parts of the northeastern regions had higher seroprevalence compared to other regions of the country. These can be used to guide the surveillance of the disease in the country.

## Introduction

Rift Valley fever (RVF) is an acute mosquito-borne viral zoonosis affecting ruminants and humans [1]. The disease is caused by Rift Valley fever virus (RVFV), which is a single-stranded RNA virus in the order *Bunyavirales* and the *Phlebovirus* genus [1]. It occurs in epizootic periods associated with heavy rainfall [2][3]. The virus was first described in detail in sheep in 1930s in the Rift Valley area of Kenya, although it may have occurred earlier [4].

In animals, the disease is characterized by high rates of abortion and neonatal mortality, primarily in sheep, goats, cattle, and camels. Humans become infected following a bite from an infected mosquito, or after close contact with acutely infected animals or their tissues [5]. Infection in humans causes a mild influenza-like syndrome in many cases (> 80 per cent), or a severe disease with haemorrhagic fever, encephalitis, or retinitis in a few cases [6].

Outbreaks of RVF were first reported in Kenya in the 1930s [4]. Since then, many epidemics have occurred in various regions including Egypt (1977), Kenya (1997–1998), Saudi Arabia (2000–2001), Yemen (2000–2001) and Kenya (2006-2007) [7–10]. In East Africa, outbreaks often occur after prolonged and heavy rains and floods [2]. Certain environmental factors such as dense vegetation, temperature conditions that favour mosquito breeding and virus replication, and presence of ruminants such as cattle, sheep and goats that can amplify transmission of the virus [11]. Areas with a flat topography, water-retaining soil types and dense bush cover are important factors for flooding and/or mosquitoes breeding [12]. The 1997–1998 RVF outbreaks in Kenya were associated with El Niño rains and floods, with consequential several human and livestock deaths [8]. In 2006–2007, Kenya, Sudan, Tanzania, and Somalia experienced a major RVF outbreak involving large numbers of humans and livestock [12].

Uganda reported an RVF outbreak after a long interval in March 2016. Historical reports of RVF in Uganda date back to 1944 when an RVF virus was first isolated from mosquitoes captured in forest in western Uganda in 1944 [13]. That isolate was later developed the Smithburn modified live virus vaccine. Another RVF occurrence was reported in 1968 in humans and livestock in Kabale district in south-western Uganda [14]. Since the Kabale 2016 outbreak, other sporadic cases have been experienced in different parts of the country. Between August 2017 and August 2018, Uganda experienced frequent RVF outbreaks in the districts of Kiboga, Kyankwanzi, Kiruhura, Isingiro, Ibanda, Lyantonde, Sembabule, Mubende, Mbarara, Kasese, Sheema, Arua, Buikwe, Lwengo and Mityana [15].

A few cross-sectional surveys have been done in the country to estimate RVFV seroprevalence in selected areas. A seroprevalence of 6.7% for anti-RVFV antibodies was reported among the domestic ruminant population in samples that were collected during a yellow fever outbreak investigation of 2010–2011 in the northern region [16]. During the 2016 outbreak in Kabale district, the seroprevalence of RVFV in southwestern Uganda was reported to be 27%, 7% and 4% and 12% in cattle, goats, sheep, and humans respectively [17]. These findings are similar to those that have been reported in Kyela and Morogoro districts, Tanzania during inter-epizootic periods where anti-RVF IgG prevalence of 29% was observed in cattle [18]. A survey that was conducted in goat farms in central Uganda in 2009 showed seroprevalences of anti-RVF IgG and VNT in sheep and goats of 9.8% and 24.3%, respectively [19].

While the various seroprevalence studies that have been implemented have illustrated the burden and risk factors for RVF in various locations, more work is still needed to map the distribution of the disease risk across the country. Such findings would enable decision makers to prioritize areas that should be targeted for surveillance and control. Although areas that are often affected by previous outbreaks are known, seroprevalence data provide a more complete picture of risk since most of the areas with endemic RVFV transmissions would be captured. RVF outbreaks are known to occur in areas with endemic RVFV transmission when conditions for their amplification are provided.

This study used data collected from a national serosurvey that involved cattle, sheep and goats to estimate RVFV seroprevalence and to map the spatial distribution of RVFV in Uganda. The study was implemented at a time when many state-of-the-art tools and procedures for mapping disease risk had become available. These include geostatistical models often founded on Bayesian inference, machine learning algorithms which have been applied extensively in species distribution modelling, and the standard hierarchical models which can be structured to account for spatial dependencies. Machine learning models (such as artificial neural networks, classification and regression trees, maximum entropy, climate envelops, and generalised linear models) suffer from statistical challenges including their inability to model various sources of uncertainty, observer biases, sampling errors, and analytical biases especially with complex spatial or spatiotemporal data. The integrated nested Laplace approximation method that was applied in this work offers more realistic and accurate predictions than standard models.

## Methods

### Study area

Uganda lies between the coordinates 29° - 35° East, and -2° - 4° north. The country’s total area is 241,038 square kilometre (sq. km) out of which 197,100 sq. km is land while 43,938 sq. km is under water. It is estimated that Uganda’s 72.1% of the land cover is agricultural land. In 2011, it was estimated that 34.3% of the land was arable, 11.3% was covered by permanent crops, 25.6% was permanent pasture, 14.5% forest cover, while about 14.3% was under other use [20].

Altitude ranges from lowest point at 621m in areas of Lake Albert to Highest point at about 5,110m on the Margherita Peak on Mountain Rwenzori above sea level. Uganda’s climate is variable and susceptible to flood and drought events which have had negative socio-economic impacts in the past [21]. The country experiences two dry seasons (December to February and June to August). Uganda’s livestock population includes 14.2 million cattle, 16 million goats, 4.5 million sheep, 47.6 million poultry and 4.1 million pigs [22].

### Study design

A farm (or household in pastoral areas) was considered as a primary unit of analysis and hence the number of farms (or households) that were required was determined using the standard sample size estimation technique for determining a population proportion. Given that the national prevalence of RVF had not been determined in the country, we use *a priori* prevalence estimate of 50% (to obtain a maximum variance and hence an estimate of the sample size). In addition, it was assumed that the study would determine RVFV seroprevalence with an error of 5%, with a confidence level of 95%. We also assumed that the livestock population was infinite and so there was no need for finite correction of the crude estimate generated. This process yielded a naïve sample size of 385 animals.

Recent studies however show that RVFV seroprevalence clusters at the herd and village levels. An intra-herd correlation coefficient of 0.3 has been determined in studies conducted in Tana River Kenya [23]. This information was used to adjust the naïve sample size by deriving the design effect, *d* using the formula:

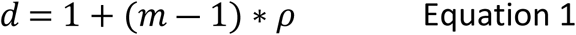

where, *m* represents mean number of animals that would be sampled per herd (assumed to be 25), *ρ* is the correlation coefficient of 0.3. A design effect of 8.2, and therefore a revised sample size of 3,157 were estimated. This implied that at least 126 herds were required for the survey.

The identification of farms/herds to sample used a multi-stage random sampling technique. First, Parishes, the smallest administrative unit, were randomly selected, followed by a selection of at least one farm in each Parish. In farms (or households) selected, at least 14 sheep or goats and 14 cattle were sampled. A herd had to have cattle, sheep and goats that were being raised together to qualify for selection.

### Data collection

#### Primary data

Four data collection teams were set up and given specific locations to sample. A team was led by a senior epidemiologist from the Ministry of Agriculture, Animal Industries and Fisheries (MAAIF), and it included laboratory scientists from the National Animal Disease Diagnostics and Epidemiology Centre (NADDEC), enumerators and animal handlers.

Once a target herd/household was identified, the team assigned temporary identification number to the household. Data on the household, herd and an animal to sample were collected using a structured form. These included administrative location and village of the farm (or household), name of the household head, herd size, and geographical coordinates (captured using hand-held GPS units in decimal degrees). The form also captured an animal identification number, age, sex, and species. Translators worked with the local people to capture data from respondents who were illiterate.

For sample collection, up to 10 ml venous blood was collected from each animal through jugular venepuncture. Injection sites were first disinfected with 70% isopropyl alcohol before the procedure was performed. The sample was collected using non-heparinized vacutainer tubes to allow for blood clotting and collection of serum. Animals from small herds were restrained within their kraals while big herds were driven to a crush for ease of handling. Blood samples were allowed to clot and later centrifuged at 3000xg for 10 minutes to harvest serum. Samples collected were kept at 4°C in the field using dry ice. They were transported to NADDEC in dry ice where they were kept at -20°C until they were analysed.

#### Secondary data

Historical data on RVF outbreaks in livestock observed between 2016 and 2021 were obtained from the Ministry of Agriculture, Animal Industry and Fisheries (MAAIF). These were to be used to determine their relative distribution with respect to RVFV seroprevalence data.

Climate and other environmental data were also obtained from various on-line databases. A total of 53 variables were included in the list. Data on their sources and spatial resolution are provided in S1 Table. The data included bioclimatic variables, soil types, altitude, pH, and vegetation indices among others, and they were included as potential predictors for RVFV.

### Laboratory screening of serum samples

At NADDEC, samples were barcoded before being stored in the sample storage freezers. All the serum samples were screened in duplicates using the Rift Valley Fever Competition Multispecies ELISA kit (ID Screen®, Grabels, France) following the manufacturer’s recommendations. The kit detects both IgG and IgM antibodies directed against the RVFV nucleoprotein. This test has been shown to have a high sensitivity (91-100%) and a high specificity (100%) in tests done by the manufacturer and an independent ring trial [24]. Before the screening commenced, two key staff from NADDEC who led the screening were invited to the International Livestock Research Institute (ILRI) in Nairobi to develop laboratory protocols for screening with ILRI staff who had used the kit before. The protocols developed included results capture spreadsheets that had in-built formulae for defining exposure categories (positive, negative, or borderline) based on optical densities obtained from an ELISA reader in accordance with manufacturer’s instructions.

### Data management and analysis

#### Data management

Both the field and the laboratory data were entered into a relational data base designed using the Census Survey and Processing System (CSPro). After cleaning, the data were exported to R (version 3.4.1) for analysis. All the animal level data including age, sex and species were captured in the database using factor variables. Age of an animal, for example, was labelled using three-level factor variable as (i) calf, lamb or kid for those animals that were still suckling, (ii) weaner for animals that had already stopped suckling but were still immature, and (iii) adults for animals that were already mature and were in breeding stages. Livestock species could be identified as ovine, caprine, or bovine.

The secondary data illustrated in S1 Table were downloaded as raster (tif) files and imported to R. These variables were scaled using the *scale*() function while they were being imported into R to enable a direct comparison of their effects. Values of these variables at sampling points were extracted using the raster extraction function that uses spatial coordinates to specify pixels where data could be extracted from.

#### Descriptive analyses

The number of herds screened, proportion of animals sampled by species, sex and age were determined by using simple descriptive statistics. The overall RVFV seroprevalence and its confidence interval was determined using the function *binconf*(). Asymptotic confidence intervals were preferred to the others that the function could provide. The estimate generated was stratified by age, sex, and species of an animal and Chi square tests used to determine whether there were significant differences between the levels of the factor variables considered.

A crude map showing the distribution of RVF outbreaks and RVFV seroprevalence across all the areas sampled. Grouping was achieved by collapsing the data at the herd level using the function *summaryBy*().

#### Cluster analysis

Sampling of multiple animals within herds, farms or households was expected to generate clustered data which could compromise the assumption of independence between observations while running statistical modelling. Moran’s *I* test was therefore used to examine whether there was significant clustering of seroprevalence data by location given that each herd shared similar geo-coordinates. The geo-coordinates that were collected during sampling were used to construct matrices that the function used to define nearest neighbours based on the function *knearneigh*().

An intra-herd correlation coefficient (ICC) was also estimated after fitting a mixed effects hierarchical logistic regression model that had a herd as the random effects. Many approaches have been proposed for estimating this parameter, including the bootstrap simulation approaches proposed by MacLeod et al. [25], but a regression approach was found to be simple and accounted for other sources of variation that had to be eliminated. The coefficient and its 95% confidence internal were generated as part post-estimation analysis.

#### Analytical modelling

##### Modelling framework

After showing that substantial spatial autocorrelation existed in the data, a hierarchical Bayesian model which applies both integrated nested Laplace approximation (INLA) and stochastic partial differential equation (SPDE) was used to effectively to account for spatial correlation and generate reliable posterior distribution of model parameters. The model was fitted to the data using the R-INLA function developed by Rue et al [26].

Two models were fitted to the data. The first had a binary outcome (*RVFV positive versus negative*), represented by a Bernoulli distribution, while the second used herd-level RVFV seroprevalence as the outcome 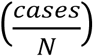, and had a Binomial distribution. The first model was used to examine both animal- and environment level factors while the second was used to develop an RVFV risk map given that it excluded animal-level variables which could not be determined in unsampled areas. For both models, the outcome was seroprevalence of RVFV, *π*_*i*_, which was linked to a linear predictor, *η*_*i*_, using a logit link. This relationship could be illustrated as:

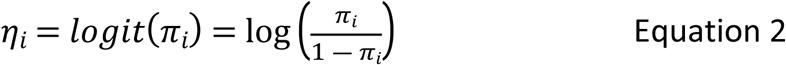

Fixed effects used in the first model included animal-level factors – sex, age, and species -- and environmental factors listed in Table S1, while the second model used environmental factors only. A general form of the model used was:

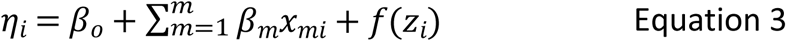

Where:

*β*_0_ is the intercept, *β*_*m*_ represent a vector of *m* coefficients (corresponding to the number of variables fitted and kept in the model), and quantifying the linear effect of covariates *x*_*m*_, and *f* (*z*_*i*_) is a function used to account for the spatial random effect.

##### Fixed effects

A combination of forward and backward variable selection procedures were used to identify and select fixed factors that could be retained in the model. A variable was considered as being significant if its 95% credible interval excluded zero. Plausible interaction terms were also examined. Similarly, linearity assumption for each for each variable was assessed by fitting quadratic functions for each variable at a time and evaluating the significance of the quadratic term.

##### Spatial effects

The spatial effect was accounted for using stochastic partial differential equations (SPDE) which is suited for modelling geostatistical data. SPDE model assumes that the random effect at each point (location) is a stochastic process with a Gaussian distribution whose mean is zero and variance estimated using Matérn variance function. A full description of this covariance structure is beyond the scope of this paper, and it has been fully described by [27]. In general, SPDE model represent an underlying continuous spatial process (Gaussian field), which spans the entire domain, using discretely indexed spatial random process. This estimation process involved the: (i) construction of a mesh, and (ii) establishment of a projector matrix to connect to the connect observed data with the mesh created.

The mesh, which aids in discretization of the Gaussian field, was constructed using the *inla*.*mesh*.2*d*() function. Uganda’s shape file, downloaded from https://www.diva-gis.org/gdata, was used to set the boundary for this function. The maximum length of the edge of the triangles that form the mesh were set to be 0.2 and 5 units within and outside the domain, respectively, and a cut-off of 0.1. The cut-off parameter defines the minimum distance that can be allowed between any two points. This would also help avoid building too many triangles around the observed locations. The mesh also allows the extension of the mesh to a region outside the target domain to avoid the boundary problems associated with SPDE model. The mesh that was constructed had 5,136 vertices (Figure 1). The significance of the SPDE model was evaluated using deviance information criterion (DIC) statistic. Models that had relatively lower DICs were preferred. Parameters of the SPDE model that may be required for future studies as priors including variance nominal, range nominal, Tau and Kappa were extracted from the marginal posterior distribution.

**Figure 1.**
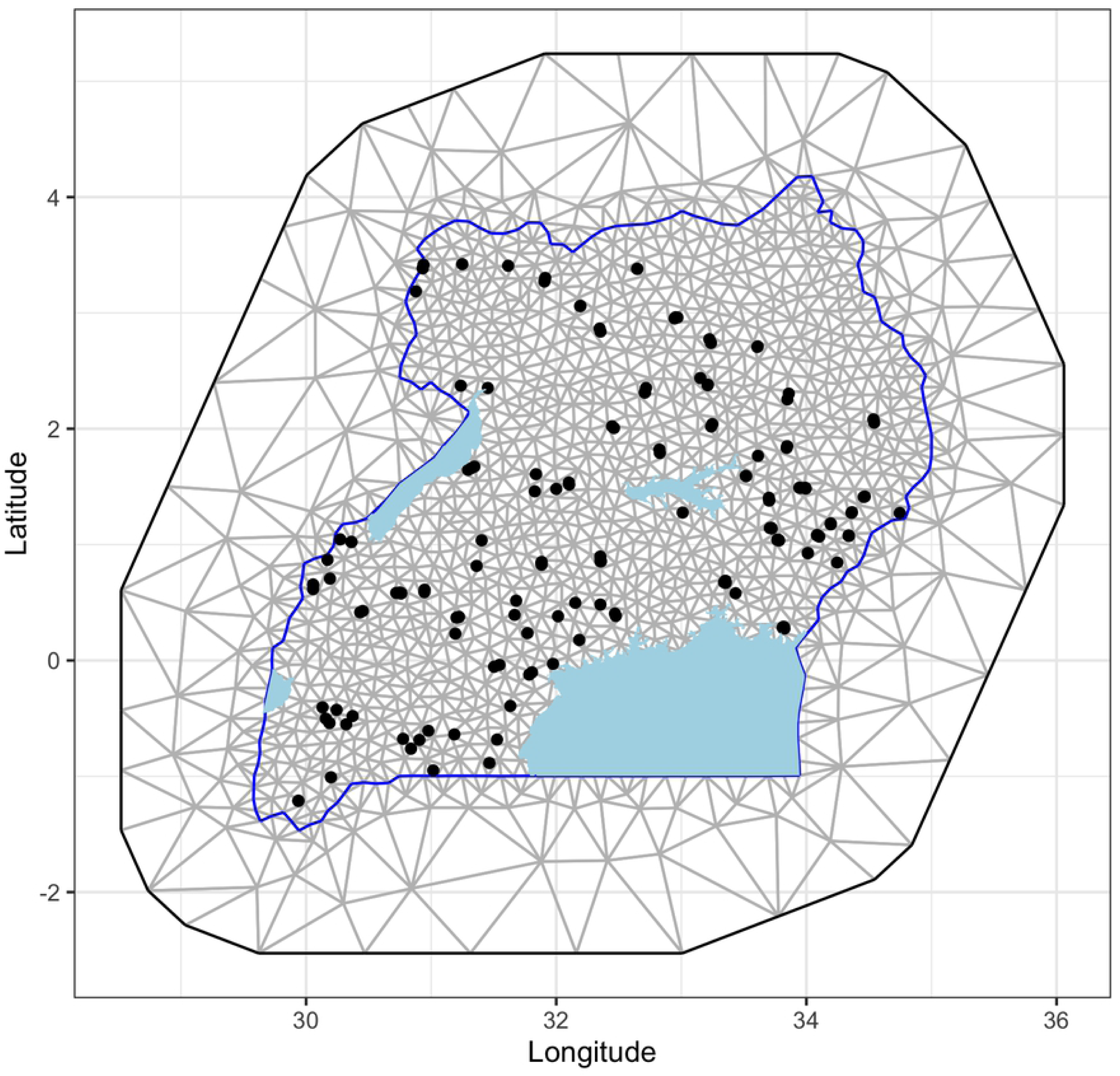
Triangulation for spatial analysis of RVFV seroprevalence in Uganda. The map indicates locations (dots) that were sampled. Uganda shape file used to demarcate the boundaries as obtained from https://www.diva-gis.org/gdata

A projector matrix was designed to link the latent field, represented by the processes modelled at the mesh vertices, to the locations of the response (coordinates obtained from the survey). The matrix was constructed using the function *inla*.*spde*.*make*.*A*(). Data stacks combining the outcomes, the predictor variables and the projector matrix was developed called into the main model command as objects.

##### Prediction of RVFV prevalence

The final version of the second model (with environmental factors alone) was used to predict RVFV seroprevalence across the domain. First, a 2km grid was generated and centroids from the grid were used to extract predictor variables. Posterior distributions of fitted values and their standard deviation were generated and mapped using *ggplot*() function.

##### Research ethics

This study obtained approval from the Uganda National Council of Science and Technology (UNCST) (Research Registration Number A603). It also received ethical approval from The International Livestock Research Institute’s Research and Ethics Committee (IREC), certificate number IREC 2017-19. Its animal sampling activities were reviewed by the institute’s Animal Use and Care Committee (IACUC). Only participants that were willing to take part in the study were recruited on informed consent. The research team practiced procedures of biosecurity and safety by donning protective gear during sample collection and analysis. All used sharps were collected in a sharps container that would later be disposed of properly at NADDEC laboratory, Entebbe. All other waste was collected in biohazard bags and incinerated at NADDEC. The clot and the used vacutainers were decontaminated with ultra chlorox bleach and ethanol 70% and incinerated properly to avoid environmental contamination.

## Results

### Descriptive statistics

A total of 3,253 animals from 175 herds were sampled. Females constituted a larger percentage (74.5%) of these animals than males (Table 1). On species distribution, cattle formed a higher majority followed by goats and sheep in that order. Most of the animals were adults; a few weaners and calves, lambs or kids were also sampled. Herd sizes ranged between 1-59 with, with a mean of 17.30 (SE=0.81).

**Table 1.** Association between animal-level factors and RVF seropositivity in Uganda.

| Factor | Level | n | Seroprevalence (95% confidence interval) | $\chi^2$; P>\|Z\| |
| --- | --- | --- | --- | --- |
| Species | Cattle | 1,692 | 15.72% (13.99 – 17.46%) | 78.59; 0.00 |
|  | Goats | 1,377 | 5.59% (4.38 – 6.81%) |  |
|  | Sheep | 184 | 10.87% (6.37 – 15.37%) |  |
| Sex | Female | 2,425 | 11.79% (10.51 – 13.08%) | 3.62; 0.06 |
|  | Male | 825 | 9.30% (7.32 – 11.28%) |  |
| Age | Adult | 1,461 | 16.70% (14.79 – 18.61%) | 36.04; 0.00 |
|  | Weaner | 838 | 8.47% (6.59 – 10.36%) |  |
|  | Calf/kid/lamb | 573 | 8.20% (5.60 – 10.45%) |  |

#### RVF seroprevalence

The overall RVF virus seroprevalence and its 95% confidence interval was 11.25% (10.17 – 12.33%). The distribution of RVF seroprevalence across the animal-level characteristics (species, sex, and age) described earlier is given in Table 1. These findings show that cattle had significantly higher RVF virus seroprevalence compared to goats, but not significantly different from that of sheep. They also show that the adult animals had significantly higher seroprevalence compared to weaners or young ones. There weren’t any differences on RVF virus seroprevalences between weaners and young (calves, kids, or lambs), or between female and males.

Figure 2 shows the distribution of sites where livestock were sampled. A colour scheme, ranging from dark blue to red, has also been used to represent RVFV seroprevalence. Of the herds sampled, 69 did not have any seropositive results, while 89 had seroprevalences of less than 30%. The rest had higher exposure levels. The map (Figure 2) also demonstrates that herds with higher RVFV seroprevalences were distributed widely across the country, specifically from the southwestern part of the country, across the central to the eastern region.

**Figure 2.**
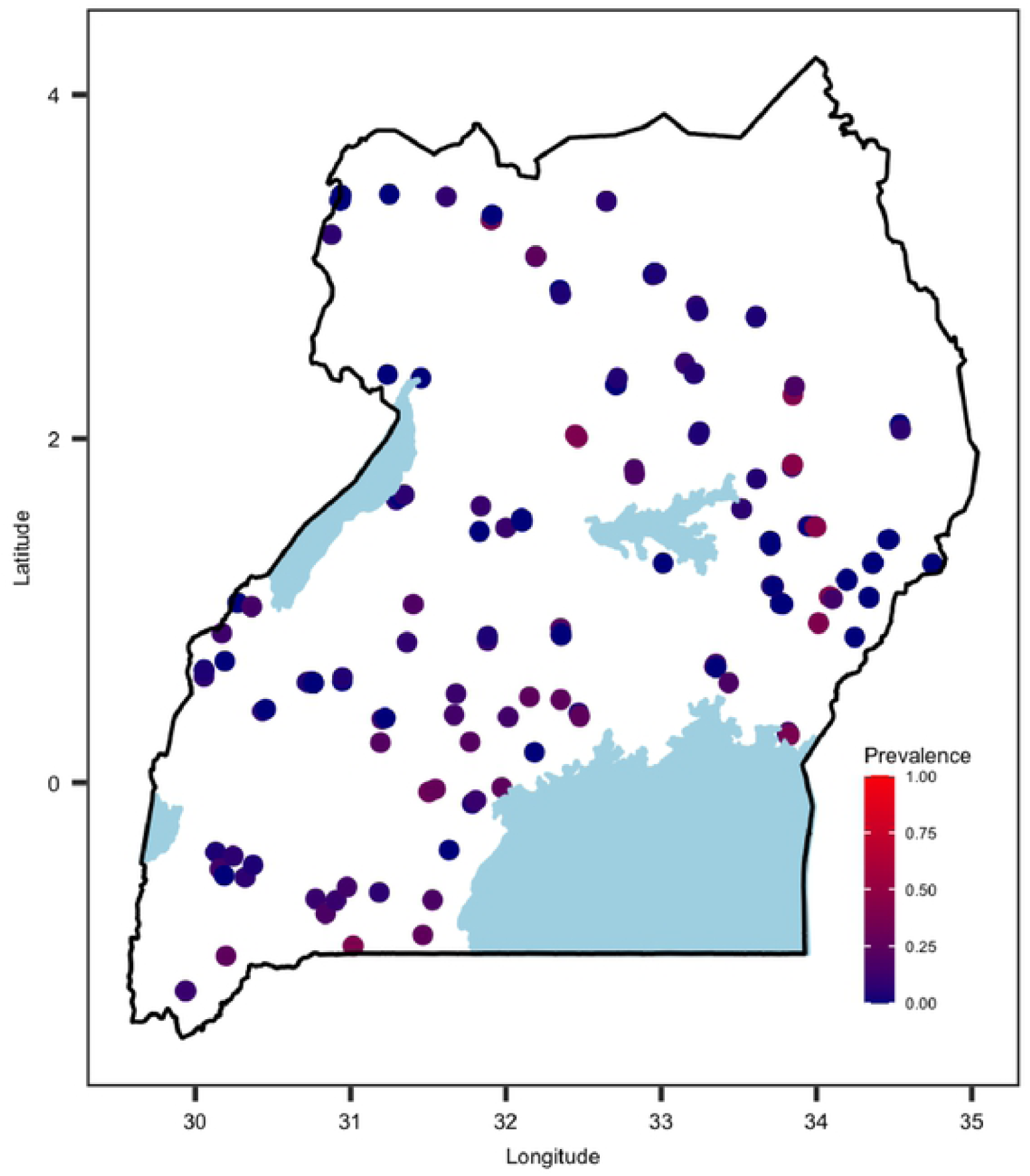
The distribution of sampling sites used for the national RVFV serosurvey and the associated levels of RVF virus seroprevalence obtained from ELISA IgG screening.

The distribution of historical RVF outbreaks observed between 2016 and 2021 are shown in Figure 3. Most of the affected areas are in the southwestern region of the country. These locations lie within, or are contiguous, to the “cattle corridor”, an ecological zone that spans diagonally across the country from southwest to northeast where rangelands are located.

**Figure 3.**
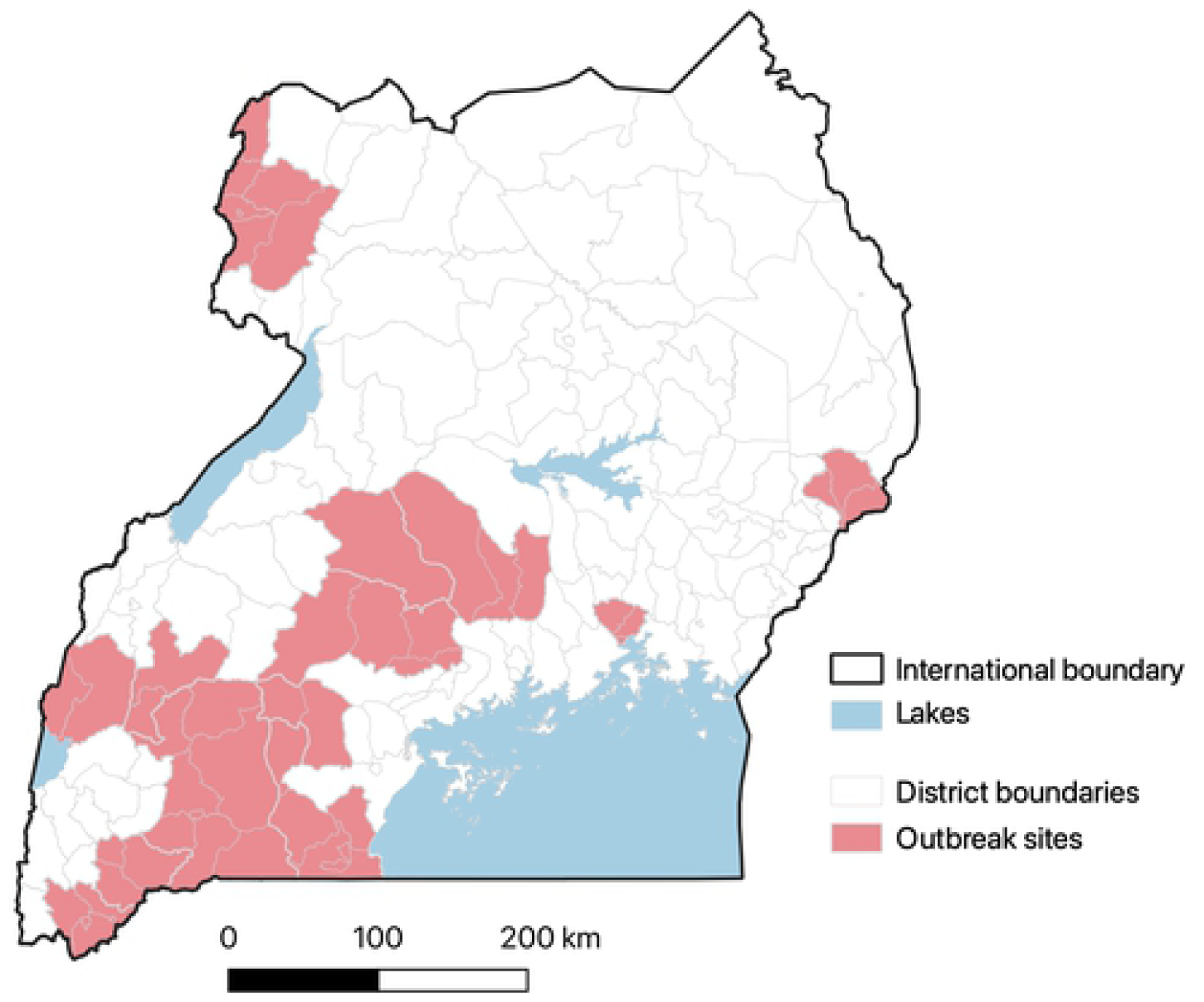
Map of Uganda showing districts that have been affected by RVF outbreaks since March 2016 when the initial outbreak in the recent years was reported. The shape file used was downloaded from https://www.diva-gis.org/gdata and outbreak data were provided by the Ministry of Agriculture, Animal Industries and Fisheries (MAAIF)

#### Cluster analysis

The Moran’s I test produced significant results (with a standard deviate of 7.60, p=0.00) suggesting that a significant spatial autocorrelation existed in the data. At the same time, an intra-herd correlation coefficient of 0.20 (95% CI: 0.04 – 0.28) was estimated.

### Results of multivariable models

Tables 2 and 3 present summary statistics of the posterior distributions of significant parameters produced by the R-INLA model. Table 2 summarises results from a model that combined animal- and environment-level variables, while Table 3 presents summarise from a model that included environment variables only. In general, including a spatial random effect improved the goodness of fit of the models. For the first model with a binary outcome, including the spatial component (SPDE model) led to a reduction in the DIC from 2006.61 to 1907.63. Similarly, the second model had its DIC declining from 657.74 to 548.60 when the spatial component was included. Five variables were significant, two of these -- age and livestock species of an animal – were animal level variables. The rest were environmental variables which included, precipitation seasonality (i.e., coefficient of variation of precipitation seasonality), cattle density, and size of areas covered by haplic planosols.

**Table 2.**
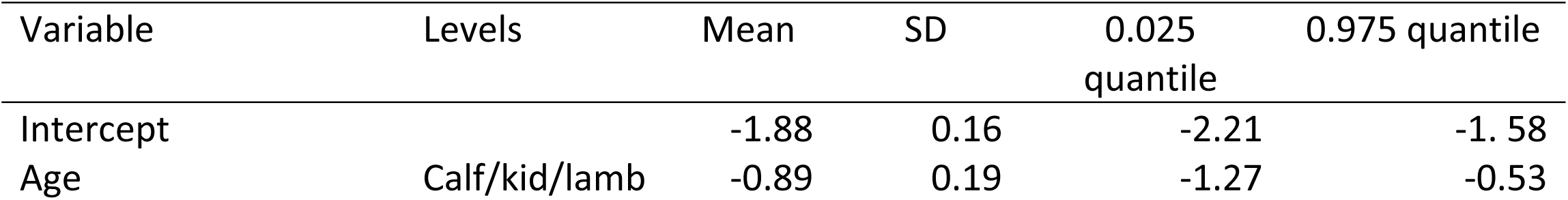
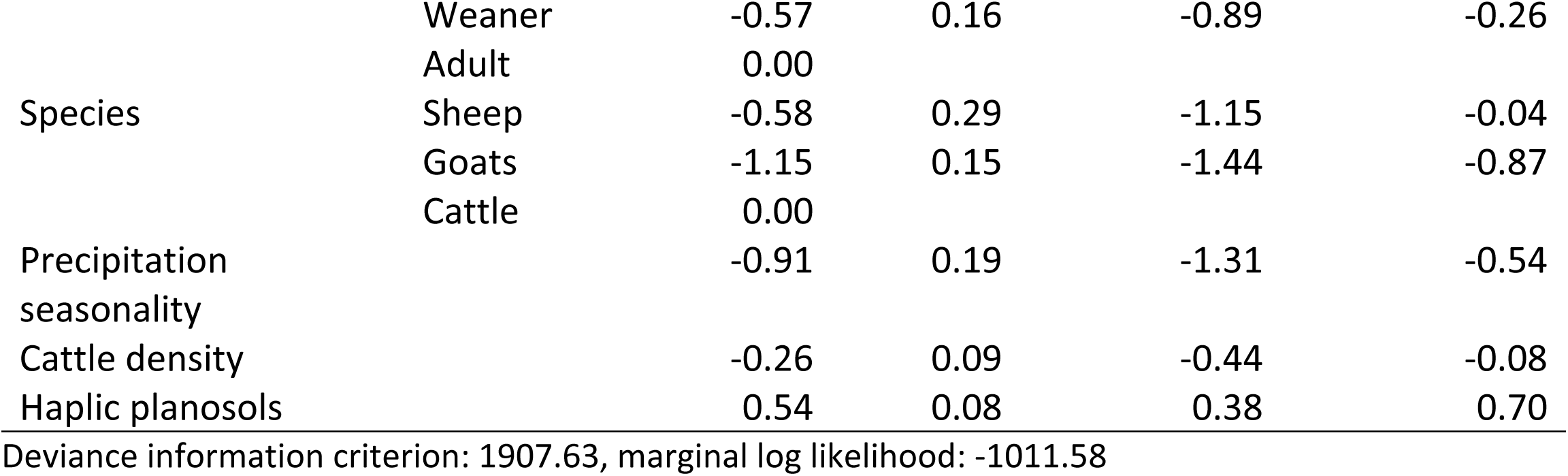
Summary statistics of posterior distributions of all the fixed effects of the hierarchical Bayesian model fitted to RVFV seroprevalence data in Uganda.

**Table 3.** Summary statistics of posterior distributions of all the fixed effects of the hierarchical Bayesian model fitted to RVFV seroprevalence data in Uganda.

| Variable | Mean | SD | 0.025 quantile | 0.975 quantile |
| --- | --- | --- | --- | --- |
| Intercept | -2.54 | 0.16 | -2.88 | -2.24 |
| Precipitation seasonality | -0.85 | 0.20 | -1.27 | -0.48 |
| Cattle density | -0.24 | 0.09 | -0.42 | -0.06 |
| Haplic planosols | 0.50 | 0.08 | 0.34 | 0.67 |

Deviance information criterion: 545.25, marginal log likelihood: -315.11

The models show that controlling for other factors, an adult animal had significantly higher log odds of exposure to RVFV than weaners or young (calves, kids, or lambs) animals (Table 2). Similarly, cattle had higher log odds of exposure to the disease compared to sheep and goats (Table 2).

Results from the second model show that an increase in coefficient of variation of precipitation seasonality was associated with a reduction on the log odds of RVFV seroprevalence (after controlling for the other two variables). Similarly, an increase in the livestock density was associated with a reduction in the log odds of RVFV seroprevalence. Conversely, a positive correlation existed between the percentage of land area that had haplic planosols with RVFV seroprevalence (Table 3). The linearity assumption for all the continuous variables (i.e., all the environmental variables) that was tested by fitting a quadratic versions of each variable was met.

#### Predicted mean RVFV seroprevalence

The posterior distribution of mean RVFV and its standard deviation are mapped in Figure 4. Three regions are predicted to have high RVFV seroprevalence – southwestern part of the country, the central region and the northeastern region. The southwestern and parts of the central regions have had RVF outbreaks in the past (Figure 5). Areas that are predicted to have high RVFV seroprevalence also have relatively higher standard deviation. The distribution of SPDE model parameters is given in Figure 6.

**Figure 4.**
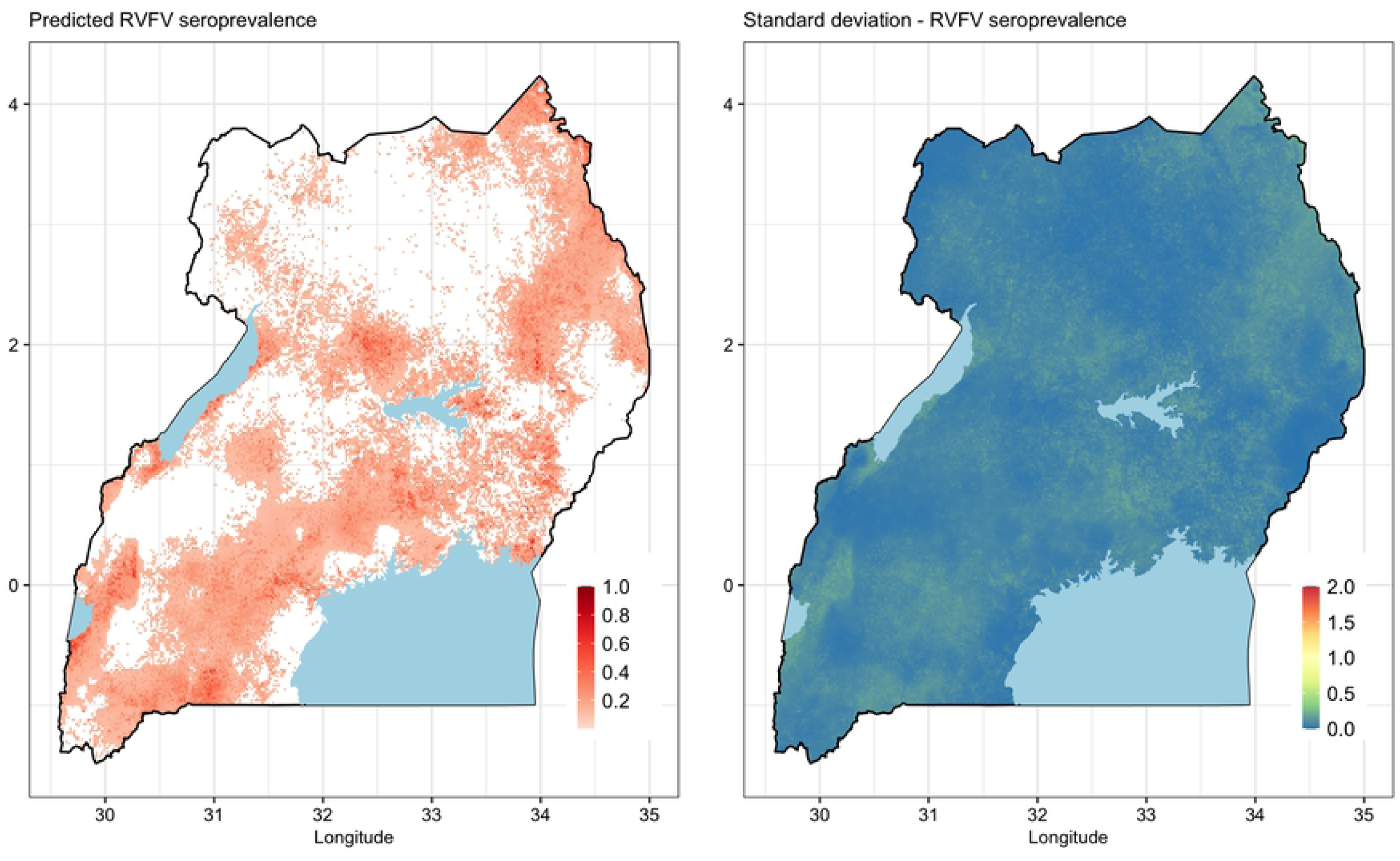
Maps of predicted posterior mean and standard deviation of RVFV seroprevalence in Uganda. Shape files used in the figure were obtained from https://www.diva-gis.org/gdata while the data were generated from the R-INLA model

**Figure 5.**
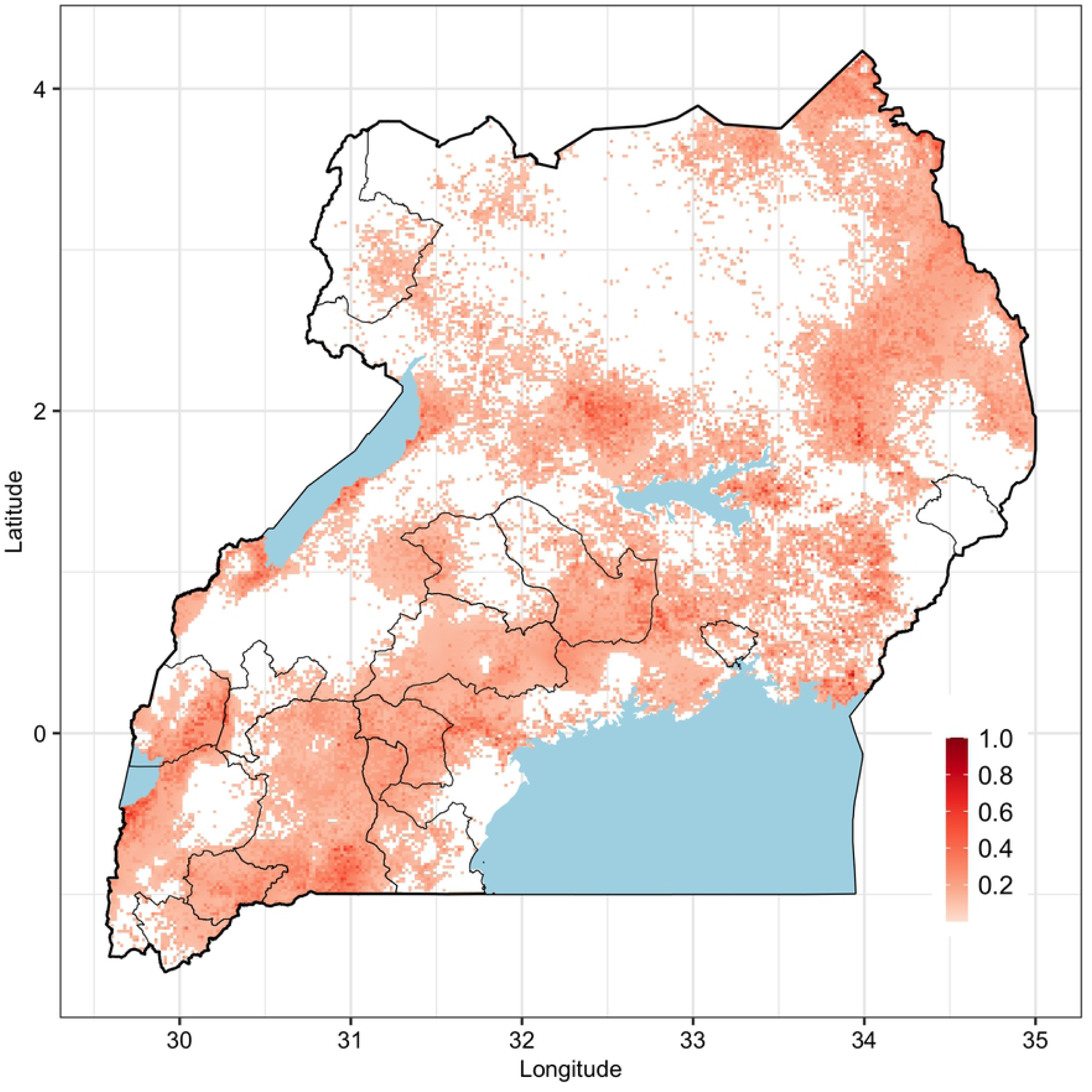
Overlaying maps of historical RVF outbreaks and predicted RVFV seroprevalence in livestock in Uganda. Shape files used in the figure were obtained from https://www.diva-gis.org/gdata. Areas affected by historical outbreaks are indicated by defined polygons in black border lines

**Figure 6.**
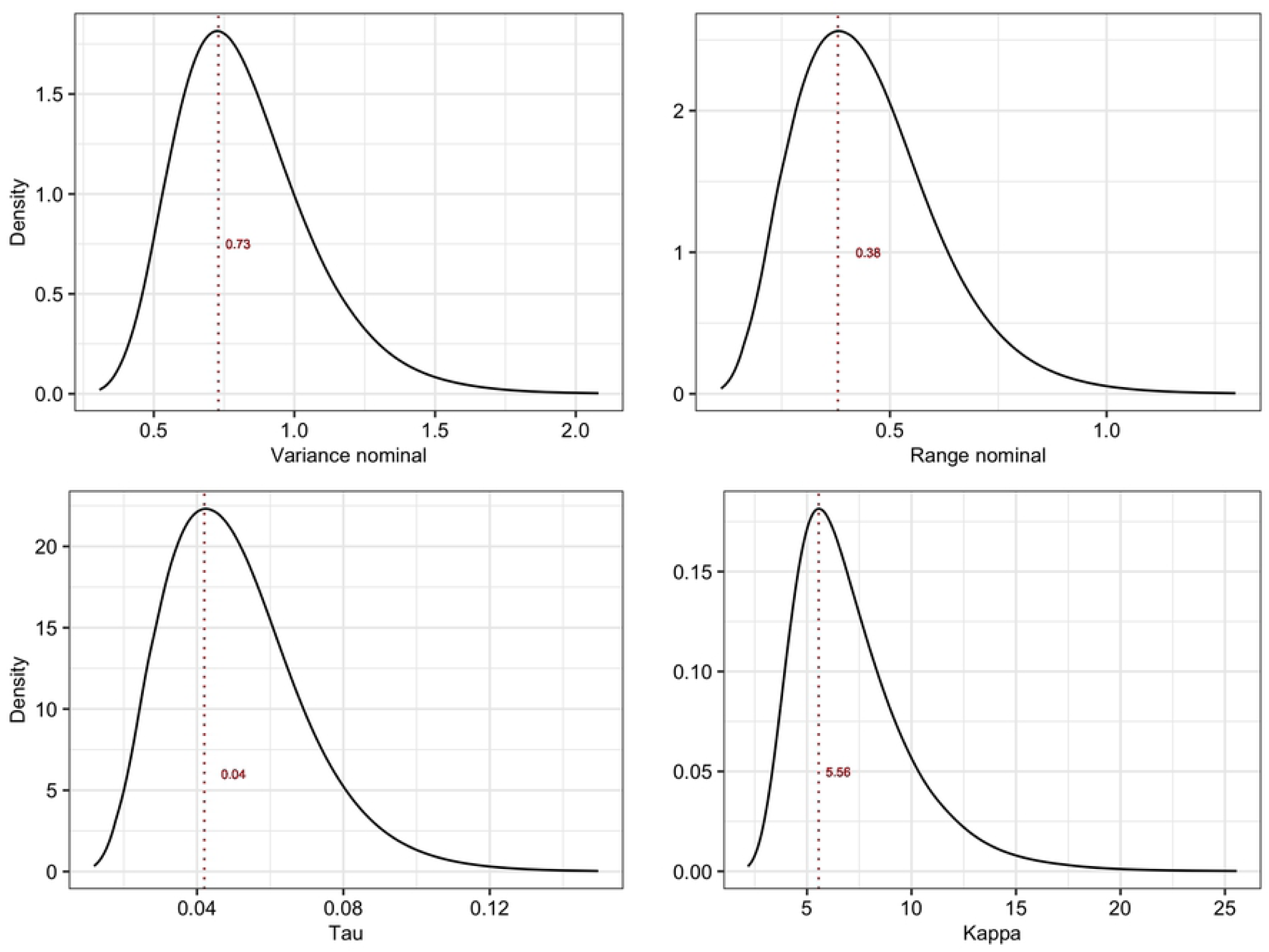
Posterior marginal distributions of the spatial parameters including variance nominal, range nominal, tau and kappa. The vertical brown dotted lines indicate mean values for each parameter.

## Discussion

This study determined spatial distribution of RVFV seroprevalence in livestock in Uganda, identified risk factors for RVFV exposure, and generated important epidemiological parameters – including intra-herd correlation coefficient, GMRF’s variance nominal, and range nominal --that could be used for advanced analyses in future. The study also compared spatial distribution of observed RVF outbreaks and RVFV seroprevalence; the significance of these observations is illustrated below.

The results from seroprevalence survey provide more reliable estimates of burden that those that have been published before given that the survey was implemented at the national level. Almost all the seroprevalence studies that have been published were implemented in the western region of the country where RVF outbreaks often occur. A recent study estimated seroprevalences of 27% in cattle, 7% in goats and 4% in sheep in Kabale district [17]. That study was however conducted shortly after the 2016 outbreak that affected the district. Similar levels of exposure in Kisoro district; seroprevalences of 20.6%, 6.8% and 3.6% were observed in cattle, sheep and goats, respectively [28]. Previously, comparable cumulative seroprevalences of 15.2% in cattle, 5.3% in sheep and 4.0% in goats in the northern, western and southwestern regions [16].

The R-INLA model used in the study identified five factors that are significantly associated with RVFV exposure. These include two animal-level factors --livestock species and age – and three environment factors – cattle density, precipitation seasonality and size of an area with planosols. As pointed out above, higher RVFV seroprevalence was observed in cattle than sheep and goats. A similar finding has been reported in other studies conducted in northern Tanzania [29] and Kenya [23]. In pastoral areas, cattle are often cover greater distances while grazing than sheep and goats to obtain adequate forage. It is therefore possible that they get exposed to more mosquito bites than the small ruminants. Cattle also have slower population turn-over rates compared to the small ruminants.

With respect to animal-level characteristics, adults had significantly higher RVFV seroprevalences compared to weaners and young animals. Similar results have been reported by many studies including those conducted in Uganda, Tanzania and Rwanda which observed a higher seroprevalence of RVF in adult animals [17,28,30,31]. RVFV anti-IgG antibodies persist for a long time in body systems, and hence the observed RVFV seroprevalences observed in older animals are likely to include cumulative exposures that have occurred over their lifetime. In addition, adults and weaners are often raised in open grazing fields that are likely to be infested with infected vectors while young animals are often tethered or kept around homesteads. The effect of sex of an animal was not significant in this study. Previous work has also produced varying results. In cross-sectional sero-surveys conducted in Kenya and Tanzania, females had higher RVF virus seroprevalences compared to males [23,30]. Studies conducted in South Africa and Rwanda, however, had different results [31,32]. It is unlikely that the sex directly influences RVF virus exposure in animals but varied management practices, including control measures and attention given to males versus females may be playing a bigger role in areas where RVF virus exposure differences are observed between sexes.

The predicted effects of the environmental variables identified in the models fitted to the data substantiate the existing knowledge on conditions that are required for persistence of RVFV transmission. These findings show that the coefficient of variation of precipitation seasonality (Bioclimatic variable No. 15 – Bio 15) is negatively associated with increased RVFV seroprevalence. This variable is a measure of the variation in monthly precipitation totals over the course of the year [33]. Water puddles or standing water masses support the development of immature mosquitoes, the biological vectors of RVFV, and areas that have stable precipitation patterns can maintain critical mosquito population levels by recharging these breeding sites. A population dynamics model that was developed for *Anopheles* spp. that transmits malaria [34] demonstrated that mosquito populations can crush out if favourable breeding conditions are not sustained. Stable precipitation patterns also sustain other critical physical conditions, such as humidity levels, that enable mosquitoes to thrive optimally.

Planosols had a strong positive association with RVFV seroprevalence in this study. These soils are predominantly found in flat waterlogged areas, and support light forests, herbs, or grasses. They have clayey properties that allow flooding when inundated. The effects of this variable should therefore be interpreted together with that of precipitation seasonality since they both contribute to the sustenance of mosquito breeding sites. A study that was conducted in Kenya following the 2006/2007 outbreak showed that RVF outbreaks that occurred in 2006/2007 were associated with four soil types – solonchaks, planosols, calcisols and solonetz [35]. That study also found out that RVF outbreaks were more common in low altitudes. While the present study identified planosols as the only significant soil type, there are important differences in the study designs used by the respective studies that need to be highlighted. The present study uses seroprevalence data which represent areas with persistent RVFV transmissions and not necessary RVF outbreaks and uses data from multiple ecological zones which may mask some relationships that may be detected in smaller regions.

Cattle density, the third environment variable that was significant, probably does not influence transmission of RVFV directly, but it may be representing other unmeasured effects in RVFV endemic areas. The correlation between cattle density and RVF occurrence has also been reported in previous studies [36].

Models fitted to the data enabled the estimation of important epidemiological parameters that are not often published, yet they are needed for quantitative analyses. The sampling of multiple animals within herds, and the determination of their geo-coordinates provided a unique opportunity for the estimation of inter-herd correlation coefficient (ICC) as well as spatial variance and range associated with GMRF process. The ICC estimate generated from the study (of 0.20) is probably the most accurate estimate given that Uganda has never vaccinated livestock against RVFV, and hence the anti-IgG seroprevalence data collected represent realistic exposure levels. An analysis that was conducted in a location in Kenya (Tana River County) estimated a much higher intra-herd correlation coefficient of 0.30 (95% CI: 0.19 – 0.44) [23] probably because that study was in a specific high-risk area of the country. ICC values are needed for determining design effect in observational studies, and they should also guide the determination of the number of herds that are required in an area for sentinel surveillance. The spatial parameters generated from the SPDE model could also be used as priors for discretising the spatial domain in in future spatial analyses. A comprehensive Bayesian inference is often limited by unavailability of informative prior parameter distributions, either for fixed or random effects. This has led to overreliance on non-informative default priors as was done in this study.

The predicted RVFV seroprevalence demonstrates areas where there is endemic transmission of RVFV in the country. The prediction of a high RVFV seroprevalence in some areas in the eastern and central parts of the country that have not reported outbreaks before (as shown in Figure 5) is probably the most significant finding given that it suggests the need to expand surveillance efforts beyond the areas that are known to have periodic outbreaks. It is likely that these areas experience covert RVF cases that go undetected by the current surveillance systems. Follow up studies are being done to verify this assumption by investigating why these areas do not experience RVF outbreaks as those with similar seroprevalences in the southwest.

The study used a cross sectional study design which, as expected, can only be used to identify risk factors, and not causes of the outcome of interest. Secondly, sampling teams were not able to visit specific areas of the country that were considered as being unsafe or accessible. The observed coverage may therefore introduce some degree of spatial selection bias.

In conclusion, the study demonstrated low-level RVF virus transmission in livestock in many parts of Uganda including those that had not reported outbreaks before. Veterinary authorities should therefore be more vigilant in their surveillance given that some of the clinical cases in these areas may have been underreported. A stronger collaboration between sectors, guided by the One Health framework, would enhance detection of any new cases especially in data from human and animal surveillance activities are integrated and analysed jointly. The study also developed an RVF risk map that would guide the authorities in the identification of RVF hotspots. Future studies should also implement longitudinal studies to better understand RVF virus circulation and maintenance in most of the high-risk areas. Further, there is need to conduct more studies including modelling association of livestock movement with RVF occurrence as well as monitoring of vector dynamics and investigation vectorial capacity and competence.

## Acknowledgements

We acknowledge Uganda’s Ministry of Agriculture, Animal Industry and Fisheries (MAAIF) for supporting the collection and laboratory analysis of the samples used. We are specifically grateful to a large team of veterinarians and laboratory technologists from the Department of Animal Health at MAAIF and the District Veterinary Officers in the districts. Those who played a key role include Flavia M. Nakanjako, Bosco C. Okuyo, Paul Lumu, Josephine Namayanja, Mary Nanfuka, Ester Nabatta, Aminah Namwabira, Eugene Arinaitwe, Beatrice Nanozi, Charity Sanyu, Franklin Mayanja, Carolyn Namatovu, Esther Nambo, and Paul Kirabo.

---

**S1 Table. List of ecological variables used for multivariable modelling**

